# Pigs are highly susceptible to but do not transmit mink-derived highly pathogenic avian influenza virus H5N1 clade 2.3.4.4b

**DOI:** 10.1101/2023.12.13.571575

**Authors:** Taeyong Kwon, Jessie D. Trujillo, Mariano Carossino, Eu Lim Lyoo, Chester D. McDowell, Konner Cool, Franco S. Matias-Ferreyra, Trushar Jeevan, Igor Morozov, Natasha N. Gaudreault, Udeni B.R. Balasuriya, Richard J. Webby, Nikolaus Osterrieder, Juergen A. Richt

## Abstract

Rapid evolution of highly pathogenic avian influenza viruses (HPAIVs) is driven by antigenic drift but also by reassortment, which might result in robust replication in and transmission to mammals. Recently, spillover of clade 2.3.4.4b HPAIV to mammals including humans, and their transmission between mammal species has been reported. This study aimed to evaluate the pathogenicity and transmissibility of a mink-derived clade 2.3.4.4b H5N1 HPAIV isolate from Spain in pigs. Experimental infection caused moderate to severe interstitial pneumonia with necrotizing bronchiolitis with high titers of virus present in the lower respiratory tract of principal-infected pigs and 100% seroconversion. Principal-infected pigs shed limited amount of virus through the nasal and oral cavities, and importantly, there was no transmission to contact sentinel pigs. Notably, critical mammalian-like mutations such as PB2-E627K and HA-Q222L emerged at low frequencies in clinical samples and tissues derived from principal-infected pigs. It is concluded that pigs are highly susceptible to infection with the mink-derived clade 2.3.4.4b H5N1 HPAIV and provide a favorable environment for HPAI viruses to acquire mammalian-like adaptations. This work is critical for further risk assessment on the ability of the newly emerging H5N1 clade 2.3.4.4b HPAIVs to cross the species barriers into mammals.

## Introduction

Influenza A viruses (IAV) are members of the *Orthomyxoviridae*, family that encompasses single-stranded, negative-sense, segmented RNA viruses. IAVs are classified into different subtypes based on the two major surface proteins: hemagglutinin (HA) and neuraminidase (NA). Wild aquatic birds are natural hosts for most IAV types, except for H17N10 and H18N11, which were first isolated from bats and seem endemic in multiple bat species ^1,2^. Highly pathogenic avian influenza (HPAI) viruses arise from the evolution of low pathogenic avian influenza (LPAI) viruses by insertion of a stretch of basic amino acids into the HA cleavage site, which enables efficient HA activation by ubiquitous cellular, furin-like proteases; they cause systemic, lethal infections in poultry ^3^.

Since the first detection of HPAI H5 subtype viruses with the isolation of the A/Goose/Guangdong/1/1996 (GsGD) H5N1 virus in southern China in 1996, this lineage has genetically evolved and spread worldwide primarily by migratory birds ^4^. Continuous evolution of the GsGD-like H5 HPAI viruses lead to the emergence of genetically distinct clades, and reassortment with other clades of HPAI viruses or local LPAI viruses has resulted in remarkable genetic diversification through the acquisition of novel gene segments ^5^. Eventually, HPAI H5N1 clade 2 viruses became dominant and their descendants have been responsible for subsequent HPAI outbreaks. One remarkable HPAI outbreak in North America happened in 2014/2015, when Asian-origin HPAI H5 clade 2.3.4.4 viruses spread to North America via the Pacific flyway (Alaska, Western Canada/USA) with migratory wild birds and underwent reassortment with local North American LPAIVs, resulting in HPAI H5 viruses with multiple NA subtypes, including H5N1, H5N2, H5N5, H5N6, and H5N8, collectively called H5Nx viruses ^6^. The geographic expansion of the H5Nx HPAI viruses via wild birds and increased virus molecular evolution resulted in the emergence and dominance of H5 subtype clade 2.3.4.4b viruses which contributed to the H5N8 outbreaks in 2016/17 and 2020/21 and the H5N1 outbreaks in 2021/22 ^7^. By 2021/22, H5N1 clade 2.3.4.4b infections had caused massive HPAI outbreaks in Europe with subsequent spread to North America via Iceland and the trans-Atlantic flyways ^8^.

The normal distribution of the receptors for avian and mammalian IAVs in the human respiratory tract usually prevents efficient direct transmission of avian influenza viruses to humans, since the respective avian-like receptor (sialic acid linked to galactose by an alpha-2,3 linkage) is found only on cells in the lower respiratory tract region ^9^. Nonetheless, in certain instances the human-animal interface allows avian influenza viruses to cross species barriers and infect humans. Human infection with avian influenza viruses occurs sporadically and results in mild to severe illness with a wide range of clinical symptoms. Specifically, it has been reported that human infections with avian H5 and H7 subtype viruses can cause severe illness in humans with case fatality rates of 52% and 39%, respectively ^10^. Human infections with HPAI H5N1 viruses have been attributed to direct exposure to infected poultry or infected wild birds. Importantly, the current epidemiological data do not provide evidence for sustained human-to-human transmission of HPAI H5 viruses ^11^. Since the beginning of 2020, human infections with clade 2.3.4.4b H5N1 have been documented in China, Spain, UK, USA, and Vietnam, and all cases were linked to close contact with infected poultry ^12^. In addition, the ability of the clade 2.3.4.4b H5N1 viruses to cross the species barrier is clearly supported by its detection in a variety of mammalian species such as terrestrial and marine mammals ^12^. In October 2022, an outbreak of the clade 2.3.4.4b H5N1 HPAI virus in farmed mink in Spain was reported, with clear evidence of sustained mink-to-mink transmission ^13^, raising concerns about the adaptation of the clade 2.2.4.4b H5N1 virus to mammals. Therefore, in order to determine the risk of the mink-derived clade 2.3.4.4b H5N1 virus isolate, this study evaluated its virulence and transmissibility in pigs, which are natural hosts for IAVs and the “mixing vessel” for the reassortment and establishment of potentially pandemic influenza viruses.

## Materials and Methods

### Cells and virus

Madin-Darby canine kidney (MDCK) cells were maintained in Dulbecco’s Modified Eagle Medium (DMEM; Corning, Manassas, VA, USA) supplemented with 5% fetal bovine serum (FBS; R&D systems, Flower Branch, GA, USA) and a 1% antibiotic-antimycotic solution (Gibco, Grand Island, NY, USA). The mink-derived clade 2.3.4.4b H5N1 isolate, A/Mink/Spain/3691-8_22VIR10586-10/2022, was kindly provided by Francesco Bonfante and Isabella Monne from the Istituto Zooprofilattico Sperimentale delle Venezie, Legnaro, Italy, and Monserrat Agüero and Azucena Sánchez from the Laboratorio Central Veterinario (LCV), Ministry of Agriculture, Fisheries and Food, Madrid, Spain, via Dr. Richard Webby from St. Jude Childrens’ Research Hospital, Memphis, TN.

### Pig infection experiment

Fifteen 4-week-old piglets were obtained from a high health status pig herd at Kansas State University. The pigs were separated into two groups: a principal-infected group and a sentinel group that were housed in separate pens. Nine pigs were challenged with a total dose of 2.2 × 10^7^ TCID_50_ in 4 mL of DMEM: 1 mL orally, 1mL intranasally, and 2 mL intratracheally (see Figure 1A). On day 2 after infection, principal-infected and sentinel pigs were co-mingled and co-housed in a single pen. Nasal swab samples were collected in 2 mL of DMEM containing 1X antibiotic/antimycotic solution at -1, 1 to 14, 17, and 21 days post-challenge (DPC), oropharyngeal swabs on days -1, 1, 3, 5, 7, 10, 14, 17, and 21 DPC. Three of the principal-infected pigs each were euthanized and necropsied at 3 and 5 DPC, and the remaining 3 principal and 6 sentinel pigs were euthanized and necropsied at 21 DPC. At necropsy, gross pathology evaluation was performed (see below) and bronchoalveolar lavage fluid (BALF) and tissue samples were collected for virological and histopathological evaluation.

**Figure 1.**
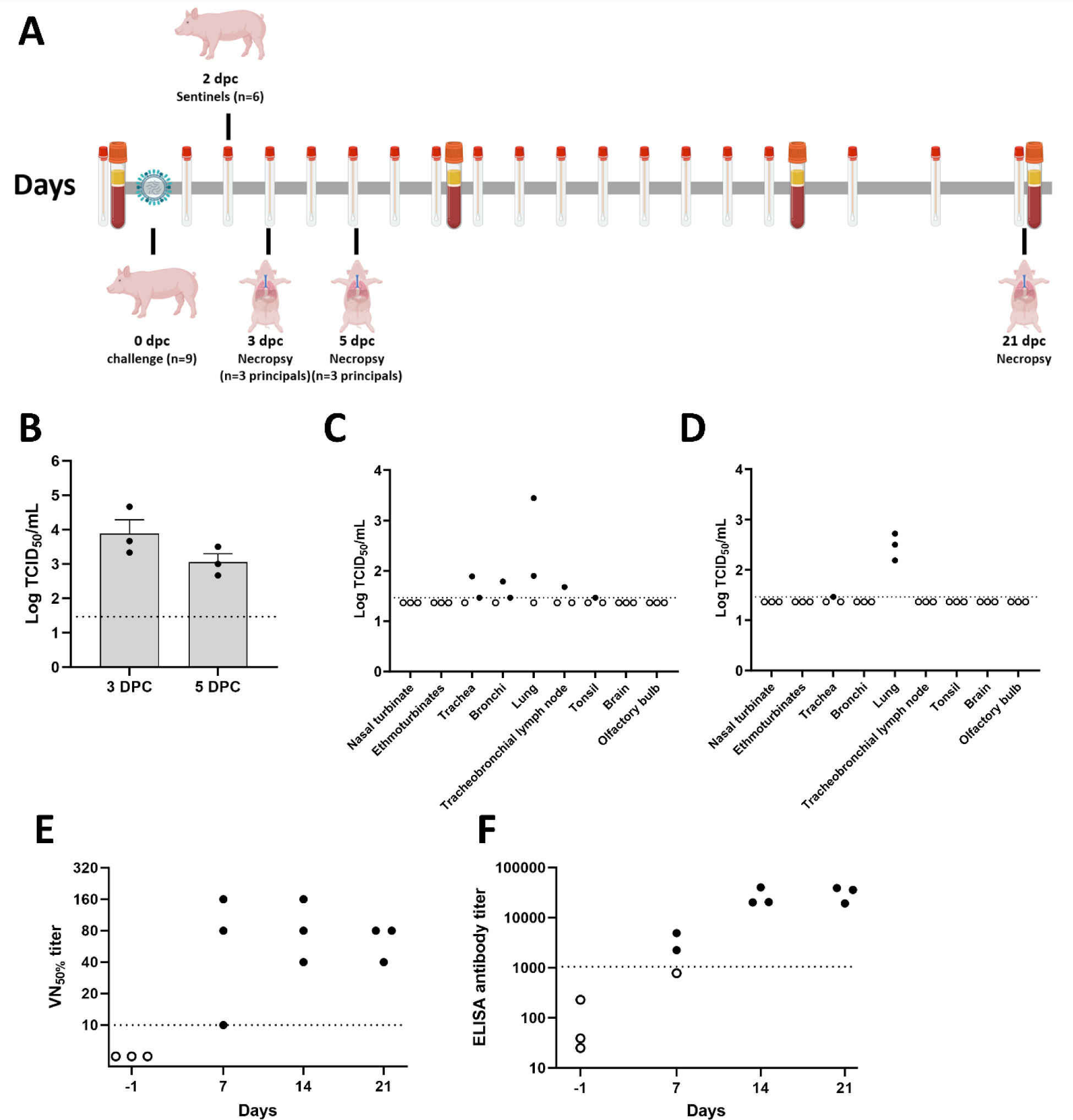
Infection with a mink-derived clade 2.3.4.4b H5N1 isolate in pigs. **A** The pigs (n=9) were orally, intra-nasally, and intra-tracheally infected with 2.2 × 10^7^ TCID_50_ of A/Mink/Spain/3691-8_22VIR10586-10/2022 and sentinel pigs (n=6) were introduced in a same pen at 2 days post-challenge (DPC). Swab and serum samples were collected throughout the study. Principal infected pigs were euthanized and necropsied at 3 and 5 DPC (n=3 per time point) and the remaining three principals and 6 sentinels were euthanized and necropsied at 21 DPC. **B** Infectious virus titers in bronchoalveolar lavage fluids at 3 and 5 DPC and **C and D** infectious virus titers in tissues at (C) 3 DPC and (D) 5 DPC. **E and F** Antibody responses were evaluated in three principal pigs at -1, 7, 14, and 21 DPC. Virus neutralization (VN) titers (E) were a reciprocal of the highest serum dilution inhibiting replication of the homologous H5N1 virus. ELISA antibody titers were calculated in a swine indirect ELISA kit (ID Screen® Influenza A Nucleoprotein Swine Indirect, Innovative Diagnostics, France) where S/P ratios were calculated from O.D. values and then converted to antibody titers according to the manufacturers’ instruction. Dash lines represent the limit of detection of each assay (the limit of detection of each assay (4.64×10^1^ TCID_50_/mL for BALF titration, 2.92×10^1^ TCID_50_/mL for tissue titration, 1:10 for VN assay, and 1053 for ELISA antibody titer). Empty circle represents negative results in each assay.

### Virus isolation

MDCK cells were used to isolate the virus from clinical samples, which were filtered through a 0.45 µm syringe filter except for BALF. Ten-fold serial dilutions of samples were prepared and transferred on confluent MDCK cells after washing with PBS. The plates were subjected to immunofluorescence assay (IFA) on day 2. Briefly, the cells were fixed with ice-cold methanol and incubated with the HB65 monoclonal antibody targeting the IAV nucleoprotein ^14^. After washing, the cells were incubated with anti-mouse conjugated with Alexa Fluor 488 (Invitrogen, Carlsbad, CA, USA). Virus titers were calculated using the Reed-Muench method ^15^.

### RNA extraction and M gene-specific RT-qPCR

Viral RNA was extracted using a magnetic bead-based automated extraction system (Taco Mini and total nucleic acid extraction kit, Gene Reach, USA) ^16^. Briefly, the swabs, BALF samples and 10% tissue homogenates were mixed with an equal volume of the RLT buffer (Qiagen, Germantown, MD, USA). Two hundreds microliter of the sample lysate and 200 µL isopropanol was added into the extraction plate. RT-qPCR was performed using the previously published primers and the modified probe (A to G at 66) in qScript XLT 1-Step RT-qPCR ToughMix (QuantaBio, Beverly, MA, USA) ^17^.

### Serology

Virus neutralization tests were performed according to a previously established protocol ^18^. Briefly, equal volumes of heat-treated serum (50 µL) and 100 TCID_50_/50 µL of virus were mixed and incubated at 37°C for 1 hour. The mixture was transferred onto pre-washed MDCK cells. On day 2, IFA was performed to determine 50% inhibition of virus growth. In addition, commercially available enzyme-linked immunosorbent assay (ELISA) kits were used to evaluate the antibody responses according to the manufacturers’ instruction: (1) ID Screen® Influenza A Nucleoprotein Swine Indirect, Innovative Diagnostics, France, (2) ID Screen® Influenza A Antibody Competition Multi-Species, Innovative Diagnostics, France, and (3) IDEXX AI MultiS-Screen Ab Test, IDEXX, USA.

### Gross pathology and histopathology

*Post-mortem* examinations were conducted at 3, 5, and 21 DPC. Macroscopic pathology was scored *in toto*, and the percentage of lung lesions calculated based on the previous published method protocols ^19^. Tissue samples were fixed in 10% neutral buffered formalin, processed, and stained with hematoxylin and eosin (H&E). Microscopic pathology was evaluated based on microanatomy of the lung using previously described scoring scale with minor modifications ^20^. Each lung section was scored from 0 to 4 scale (0= absent, 1=minimal, 2=mild, 3=moderate, and 4=severe) for six separate criteria typically associated with influenza A virus infections in pigs: (i) epithelial necrosis, attenuation or disruption; (ii) airway exudate; (iii) percentage of airways with inflammation; (iv) peribronchiolar and perivascular lymphocytic inflammation; (v) alveolar exudate; (vi) alveolar septal inflammation. For scoring the percentage of airways with inflammation (iii): 0%=0, up to 10% involvement =1, 10–39% involvement =2, 40–69% involvement =3 and, greater than 70% involvement =4.

### Influenza A virus-specific immunohistochemistry (IHC)

Immunohistochemistry (IHC) for detection of Influenza virus A H5N1 nucleoprotein (NP) antigen was performed on the automated BOND RXm platform and the Polymer Refine Red Detection kit (Leica Biosystems, Buffalo Grove, IL). Following automated deparaffinization, four-micron formalin-fixed, paraffin-embedded tissue sections on positively charged Superfrost® Plus slides (VWR, Radnor, PA) were subjected to automated heat-induced epitope retrieval (HIER) using a ready-to-use EDTA-based retrieval solution (pH 9.0, Leica Biosystems) at 100 °C for 20 min. Subsequently, tissue sections were incubated with the primary antibody (rabbit polyclonal anti-Influenza A virus NP [ThermoFisher Scientific, PA5-32242] diluted 1:500 in Primary Antibody diluent [Leica Biosystems]) for 30 min at ambient temperature followed by a polymer-labeled goat anti-rabbit IgG coupled with alkaline phosphatase (30 min). Fast Red was used as the chromogen (15 min), and counterstaining was performed with hematoxylin for 5 min. Slides were dried in a 60 °C oven for 30 min and mounted with a permanent mounting medium (Micromount®, Leica Biosystems). Lung sections from a pig experimentally infected with swine influenza virus A/swine/Texas/4199-2/1998 H3N2 were used as a positive assay control.

### Influenza A virus H5N1 mink-specific RNAscope® in situ hybridization

For RNAscope® in situ hybridization (ISH), an anti-sense probe targeting the NP and the hemagglutinin (HA [H5 subtype]) of influenza virus A/Mink/Spain/3691-8_22VIR10586-10/2022 clade 2.3.4.4b (GISAID # EPI_ISL_15878539) were designed (Advanced Cell Diagnostics (ACD), Newark, CA, USA). Sections of formalin-fixed paraffin-embedded tissues were generated as indicated above, and the RNAscope® ISH assay was performed using the RNAscope 2.5 LSx Reagent Kit (ACD) on the automated BOND RXm platform (Leica Biosystems, Buffalo Grove, IL, USA). Following automated baking and deparaffinization, tissue sections were subjected to heat-induced epitope retrieval (HIER) using an EDTA-based solution (pH 9.0; Leica Biosystems) at 100°C for 15 min, protease digestion using the RNAscope® 2.5 LSx Protease for 15 min at 40 °C, and incubation with a ready-to-use hydrogen peroxide solution for 10 min at room temperature. Slides were incubated with each probe mixture for 2 h at 40 °C, and the signal was amplified using a specific set of amplifiers (AMP1 through AMP6) as recommended by the manufacturer. The signal was detected using a Fast-Red solution for 10 min at room temperature. Slides were counterstained with a ready-to-use hematoxylin for 5 min, followed by five washes with 1X BOND Wash Solution (Leica Biosystems). Slides were finally rinsed in deionized water, dried in a 60 °C oven for 30 min, and mounted with Ecomount® (Biocare, Concord, CA, USA). Lung sections from a pig experimentally infected with swine influenza virus A/swine/Texas/4199-2/1998 H3N2 were used as a positive assay control for the NP-specific probe, while lung sections from this study were used in conjunction with the former to validate the HA-specific probe.

### Next-generation sequencing (NGS)

The whole genome sequence of the mink-derived H5N1 clade 2.3.4.4b virus was determined using the Illumina MiSeq sequencing platform (Illumina, San Diego, CA, USA). Briefly, viral RNA was extracted from the challenge virus inoculum and virus-positive clinical samples using the QIAamp viral RNA mini kit (Qiagen, Germantown, MD, USA) according to the manufacturer’s instructions. Viral gene segments were amplified using SuperScript™ III One-Step RT-PCR System with Platinum™ Taq DNA Polymerase (Thermo Fisher Scientific, Waltham, WA, USA) with the Opti and Uni universal influenza primer sets ^21,22^ or with SuperScript™ IV First-Strand Synthesis System and Platinum™ SuperFi™ DNA Polymerase (Thermo Fisher Scientific, Waltham, WA, USA) with previously published segment specific primer sets^23^. All samples were normalized to 100−500 ng of DNA prior to library preparations. Sequencing libraries were prepared using the Illumina DNA Prep kit (Illumina, San Diego, CA). Libraries were sequenced using pair-end chemistry on the Illumina MiSeq platform with the Miseq v2 Reagent kit (300 cycles). Sequencing reads were demultiplexed and parsed into individual FASTQ files and imported into CLC Genomics Workbench version 22.0.1 (Qiagen, Germantown, MD, USA) for analysis. Reads were trimmed to remove primer sequences and filtered to remove low quality and short reads. The trimmed reads were mapped to the reference sequences (GISAID accession numbers: EPI2220590 to EPI2220597). Following read mapping, all samples were run through the low frequency variant caller module within CLC Genomic Workbench with a frequency cutoff greater than 2%.

## Results

### Infection of pigs with the mink-derived clade 2.3.4.4b H5N1

This study aimed to elucidate the pathogenicity and transmissibility of the mink-derived clade 2.3.4.4b H5N1 virus in pigs. At 24 hours post-challenge, all principal-infected pigs became lethargic, and five of nine pigs were febrile with average temperatures of 41 °C. However, the average temperature decreased to 39.5 °C at 2 DPC and pigs were healthy with no obvious clinical signs throughout the rest of the observation period (Supplementary Figure 1).

Infectious virus was isolated from nasal (6/9 animals positive) and oropharyngeal swabs (4/9 positive) from principal infected pigs and the titers ranged from 4.46 × 10^1^ to 1 × 10^3^ TCID_50_/mL at 1 DPC (Table 1). At 3 DPC, one nasal and one oropharyngeal swab were virus positive, with titers of 2.15 × 10^3^ and 4.64 × 10^2^ TCID_50_/mL, respectively. Infectious virus was still present in the nasal swab at 5 DPC (2.15 × 10^2^ TCID_50_/mL). RT-qPCR results showed that all principal-infected animals were positive on at least one time point from 1–5 DPC in either the nasal or oropharyngeal swabs (Supplementary Figure 2A and 2B). All swabs collected from sentinel pigs were negative by virus isolation and RT-qPCR throughout the entire observation period.

**Table 1.** Virus shedding in pigs infected with the mink-derived clade 2.3.4.4b H5N1 virus.

| Days post-challenge (DPC) |  |  | 1 | 2 | 3 | 4 | 5 | 6 to 21<sup>a</sup> |
| --- | --- | --- | --- | --- | --- | --- | --- | --- |
| Nasal swab | Principal | # of positive / # of animals | 6/9 | 0/9 | 1/9 | 0/6 | 1/6 | 0/3 |
|  |  | Virus titer (TCID<sub>50</sub>/mL) | 4.6×10<sup>1</sup> to 1×10<sup>3</sup> | Neg | 2.15×10<sup>3</sup> | Neg | 2.15×10<sup>2</sup> | Neg |
|  | Sentinel | # of positive / # of animals | 0/6 | 0/6 | 0/6 | 0/6 | 0/6 | 0/6 |
| Oropharyngeal swab | Principal | # of positive / # of animals | 4/9 | N/A<sup>b</sup> | 1/9 | N/A<sup>b</sup> | 0/6 | 0/3 |
|  |  | Virus titer (TCID<sub>50</sub>/mL) | 1×10<sup>2</sup> to 4.6×10<sup>2</sup> | N/A<sup>b</sup> | 4.6×10<sup>2</sup> | N/A<sup>b</sup> | Neg | Neg |
|  | Sentinel | # of positive / # of animals | 0/6 | N/A<sup>b</sup> | 0/6 | N/A<sup>b</sup> | 0/6 | 0/6 |

<sup>a</sup>Nasal swabs were collected at 6 to 14, 17 and 21 DPC and oropharyngeal swabs were at 7, 10, 14, 17, and 21 DPC. <sup>b</sup>N/A: Samples were not collected.

To determine virus replication and distribution in tissues, virus titers were determined in BALF and various tissue samples. All BALF samples at 3 and 5 DPC were positive with virus titers of 2.15 × 10^3^ to 4.64 × 10^4^ TCID_50_/mL at 3 DPC and 4.64 × 10^2^ to 3.16 × 10^3^ TCID_50_/mL at 5 DPC (Figure 1B). Infectious virus could also be isolated from tracheas (3/6), bronchi (2/6), lung tissues (5/6), tracheobronchial lymph nodes (1/6), and tonsils (1/6) of six principal-infected pigs that were euthanized at 3 (n=3 pigs) and 5 (n=3 pigs) DPC (Figure 1C and 1D). Interestingly, viral RNA was detectable in the ethmoturbinates of all three animals sacrificed at 3 DPC (Supplementary Figure 2C). In addition, viral RNA was present in the nasal turbinate of one pig at 3 DPC; this pig shed infectious virus from the nasal cavity at 1 and 3 DPC. In contrast, the brain and olfactory bulb at 3 and 5 DPC were negative as assessed by virus isolation and RT-qPCR (Supplementary Figure 2C and 2D). As expected, BALF and tissue samples from both principal-infected and sentinel animals at 21 DPC were negative by both virus isolation and RT-qPCR.

To determine the antibody responses in principal-infected and sentinel pigs, virus neutralization tests (VNTs) and enzyme-link immunosorbent assays (ELISAs) were performed. Neutralizing antibodies were detectable in all three principal-infected animals starting at 7 DPC and all principals were sero-positive at 14 and 21 DPC (Figure 1E), with neutralization titers ranging from 1:10 to 1:160. The NP-based indirect ELISA revealed that two out of three principal-infected animals seroconverted at 7 DPC, with the levels of antibodies in all three principal-infected pigs against NP increasing by 14 and 21 DPC (Figure 1F). Two different NP-based competitive ELISAs confirmed the seroconversion of the three principal-infected pigs at the 14 and 21 DPC (Supplementary Figure 3A and 3B).

### Macroscopic and microscopic pathology

At 3 DPC, moderate to severe, multifocal to coalescing interstitial pneumonia with marked congestion and edema affected all lobes were noted in 2/3 euthanized animals, while the remaining animal had mild, multifocal pneumonia with congestion predominately the right lung and left caudal lung lobes (Supplementary Figure 4A–4C). Pulmonary lesions consisted of moderate to severe non-suppurative interstitial pneumonia with necrotizing bronchitis/bronchiolitis affecting multiple bronchopulmonary segments, predominantly the pulmonary parenchyma adjacent to terminal bronchioles. Affected bronchopulmonary segments were characterized by degeneration, attenuation and necrosis of the bronchiolar epithelium and intraluminal cell debris and degenerate neutrophils, as well as peribronchiolar and perivascular histiocytic, neutrophilic and lymphocytic inflammation locally extending to and expanding alveolar septa. Smaller bronchi and bronchioles were most commonly affected, and necrotizing alveolitis was characterized by alveolar spaces that contain inflammatory cells, occasional fibrin and necrotic cell debris; the epithelial lining was segmentally to partially denuded or lined by swollen degenerate epithelium (Figure 3A). These microscopic changes correlated with intracytoplasmic and/or intranuclear viral NP antigen frequently within bronchial/bronchiolar epithelial cells, and less frequently within pneumocytes and alveolar macrophages (Figure 3B). Viral NP (as determined by NP-specific IHC) and H5-specific M or HA RNA (as determined by RNAscope® ISH) were predominantly detected within airway epithelia and intraluminal necrotic cell debris, and sporadically in inflammatory cells within the bronchiolar lamina propria (Figure 4A and 4B). In 1/3 pigs (#839), there was localized suppurative and erosive rhinitis affecting the olfactory region (Supplementary Figure 5A). Within the affected region, viral antigen was detected in sporadic foci of the olfactory neuroepithelium (Supplementary Figure 5B). Larger airways (segmental bronchi and trachea) were affected by mild, multifocal neutrophilic to lymphohistiocytic bronchitis and tracheitis, with frequent bronchial but rare tracheal epithelial cells containing viral antigen (Supplementary Figure 5C and 5D).

**Figure 2.**
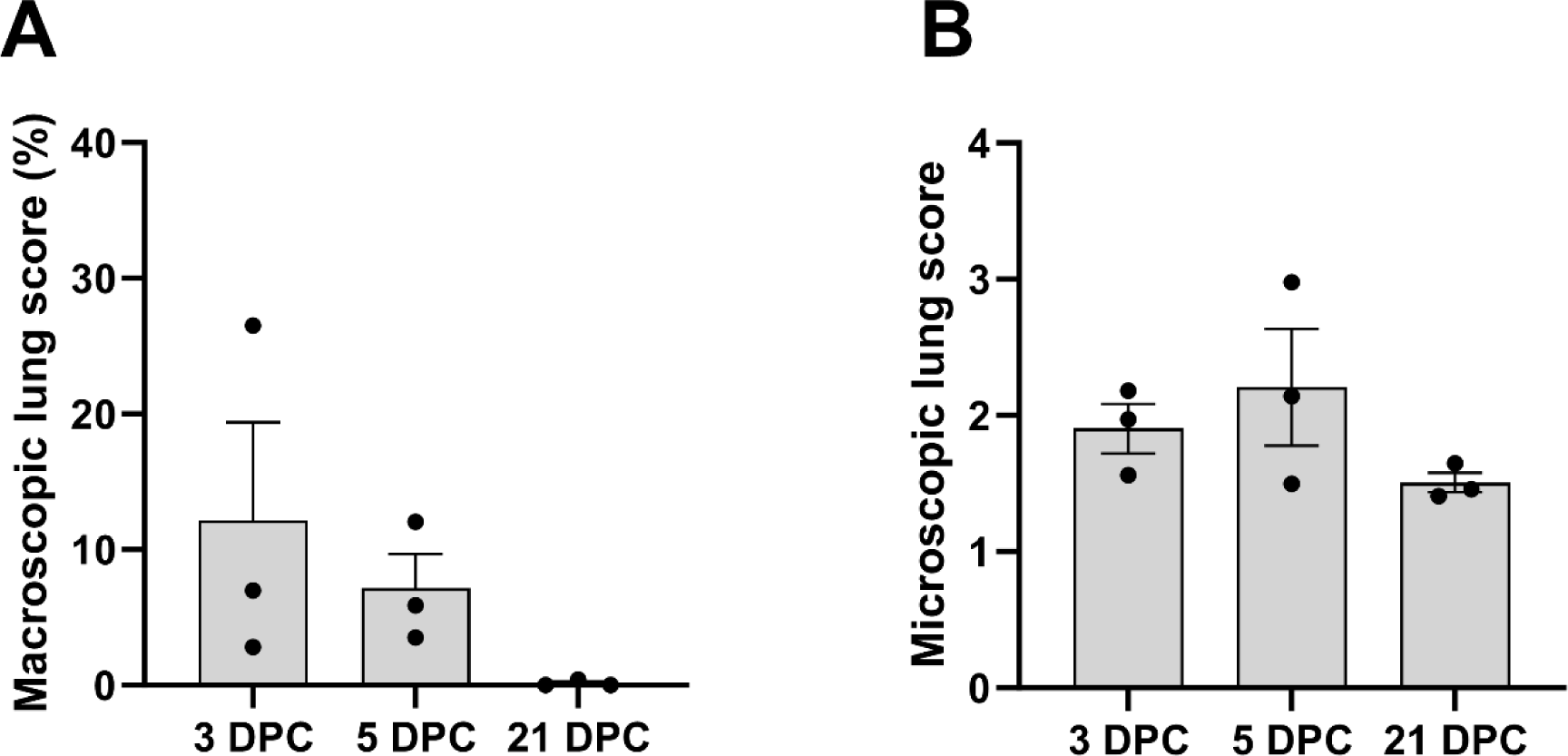
Macroscopic and microscopic lung scores in a mink-derived clade 2.3.4.4b H5N1 virus-infected pigs. Infected pigs were euthanized at 3, 5, and 21 DPC for pathological evaluation. Macroscopic lung scores were determined by the percentage of the surface area showing IAV-induced pneumonia. Microscopic scores were calculated by the average of six different criteria (see Materials and Methods)

**Figure 3.**
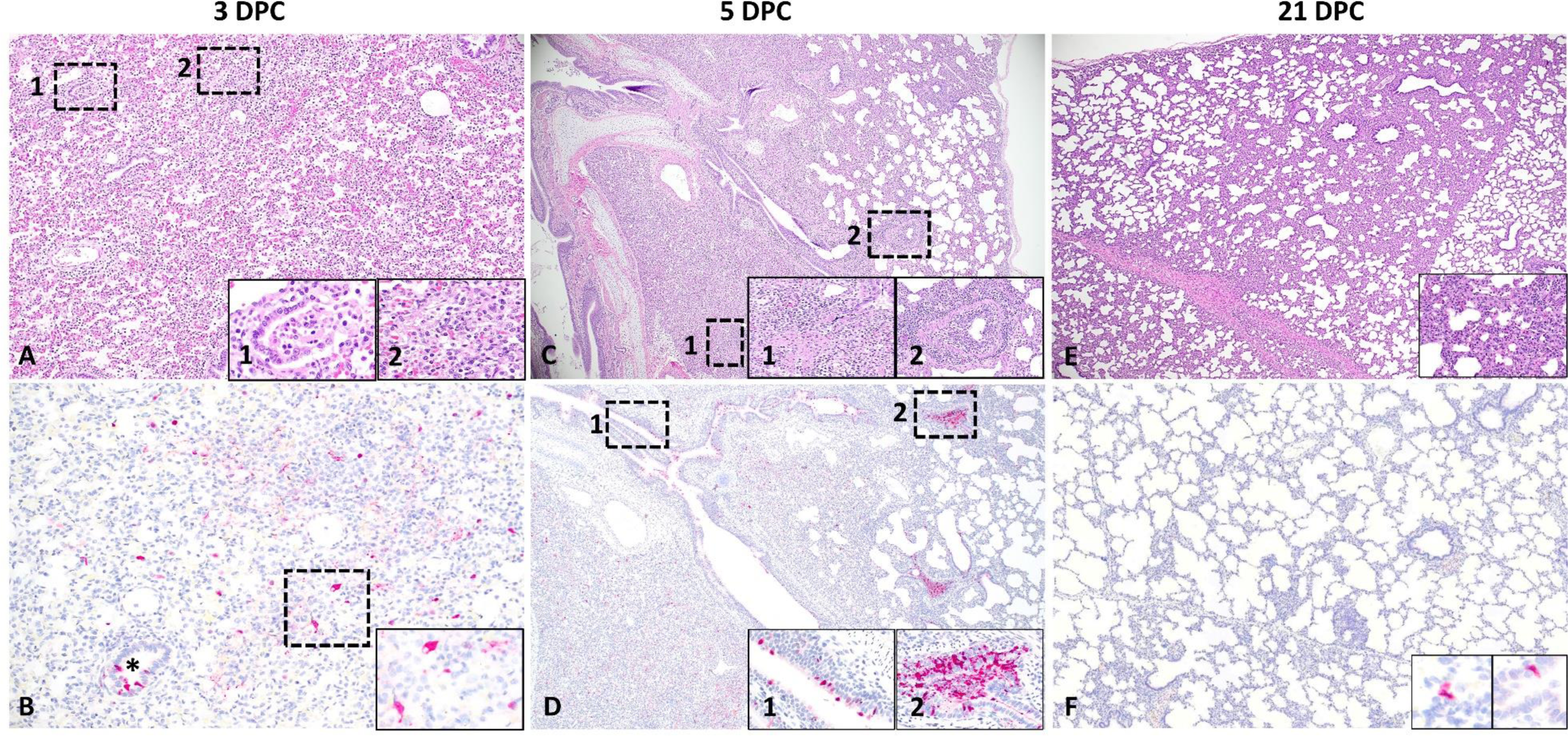
Histological features and tropism of mink-derived H5N1 clade 2.3.4.4b influenza virus in pigs. Histologically, pulmonary lesions were characterized by necrotizing to lymphohistiocytic bronchointerstitial pneumonia. At 3 DPC (A and B), multifocal bronchopulmonary segments were characterized by bronchiolar epithelial cell necrosis (A, inset 1) and associated areas of pulmonary parenchyma with expanded alveolar septa infiltrated by lymphocytes and histiocytes and alveolar epithelial necrosis (A, inset 2). Influenza A virus NP intracytoplasmic antigen was detected within bronchiolar epithelial cells (B, asterisk), pneumocytes and alveolar macrophages (B, inset). Intranuclear viral antigen was typically detected within infected epithelial cells. Similar but more extensive and severe histological changes were noted at 5 DPC (C and D), with similar alveolar (C, inset 1) and bronchiolar (C, inset 2) alterations. Influenza A virus NP antigen was more abundant but had a similar cellular distribution compared to 3 DPC (D and insets 1 and 2). At 21 DPC, the airway epithelia had mostly repaired but multifocal areas of interstitial inflammation persisted (E, inset). Only sporadic epithelial cells contained viral antigen at this timepoint (F and insets).

**Figure 4.**
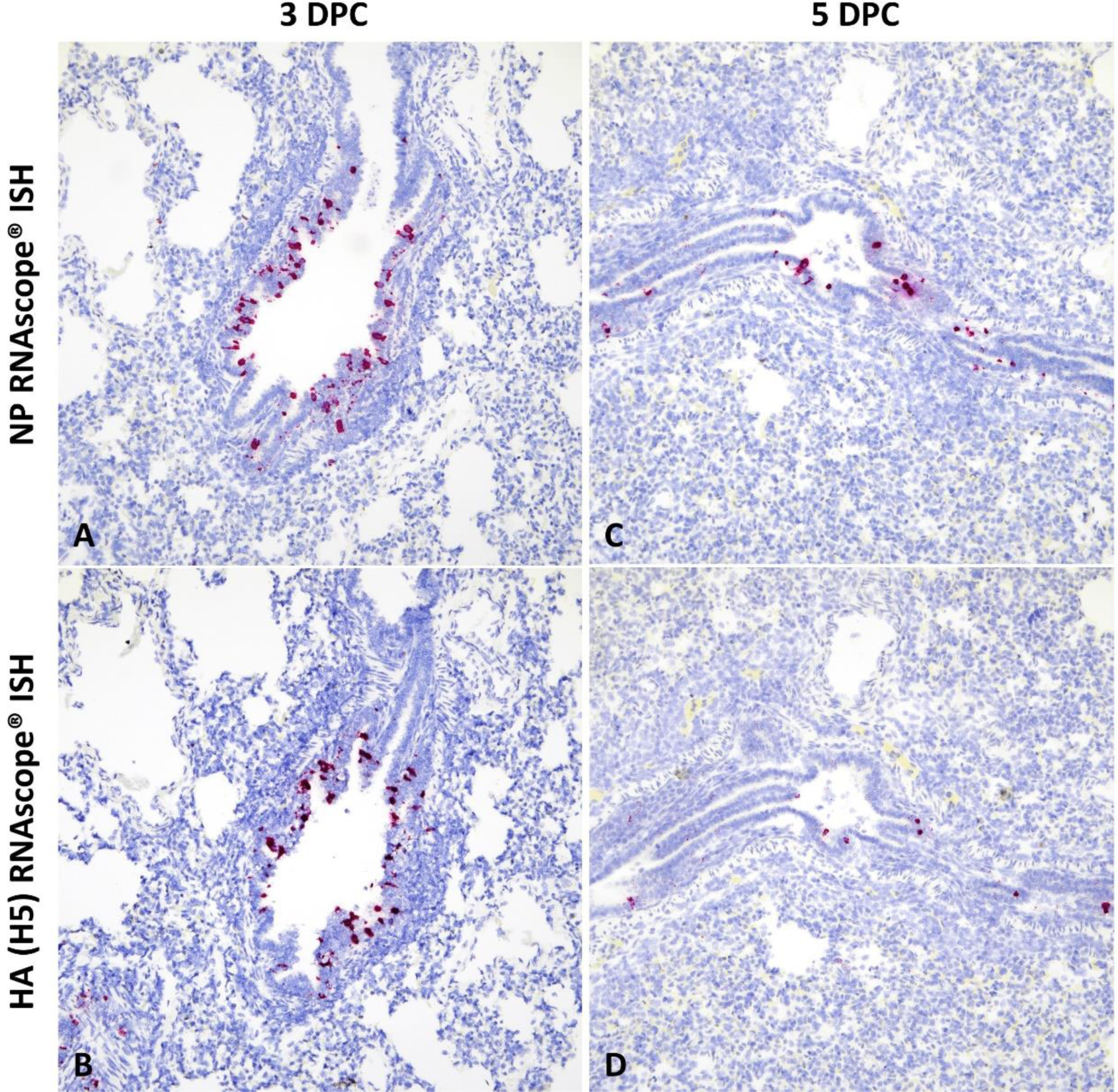
Detection of mink-derived H5N1 clade 2.3.4.4b influenza virus RNA via in situ hybridization (ISH). Probes specific to the NP and HA (H5) genes were designed. Viral RNA was predominantly detected within bronchiolar epithelial cells and intraluminal cellular debris at 3 (A and B) and 5 DPC (C and D). RNAscope® ISH, Fast Red, total magnification 200X.

By 5 DPC, multifocal to coalescing areas of pulmonary consolidation were observed on all pigs (Supplementary Figure 4D–4F). In 2/3 pigs, focally extensive pneumonia was accompanied by large air-filled bullae formation in the left dorsal lobes. Microscopically, alterations were similar to but more extensive than those at 3 DPC with intense inflammation affecting larger regions of the pulmonary parenchyma, with frequent bronchial/bronchiolar involvement and locally extensive alveolar collapse and consolidation (Figure 3C). Viral NP antigen and, less frequently, NP- and HA-specific RNA were detected in alveolar macrophages and intraluminal cell debris (Figure 3D and 4C–4D). In addition, numerous individuals and small clusters of epithelia on medium to large airways contain viral antigen. Microscopic changes in the trachea were mild and characterized by neutrophilic and lymphohistiocytic tracheitis with sporadic epithelial cell degeneration/necrosis and rare intracytoplasmic viral antigen (Supplementary Figure 5E and 5F). Multifocal, mild suppurative to erosive rhinitis affecting rostral turbinates was identified, although in lieu of intralesional viral antigen. On 21 DPC, multiple foci of mild lymphohistiocytic interstitial pneumonia were noted, accompanied by bronchiolar epithelial hyperplasia (repair) with very rare bronchiolar epithelial cells or proprial mononuclear cells containing viral antigen (Figure 3E and 3F).

Interestingly, Influenza A virus NP-positive antigen presenting cells were common within tracheobronchial lymph nodes, most abundant at 3 DPC (with an average of 10-20 cells per 200X field) and gradually reducing in numbers through 21 DPC (Supplementary Figure 6). No other significant gross or histologic lesions, or viral antigen were identified in other organs examined.

### Emergence of mammalian-like mutations in H5N1-infected pigs

The whole-length genome of the H5N1 virus inoculum, two nasal swabs, five oropharyngeal swabs and six BALF samples were genetically characterized to determine genetic evolution of the H5N1 virus after replication for up to 5 days in pigs. Our results show that the original mammalian-like PB2 T271A mutation present in the inoculum remained stable with a frequency below 2% with no evidence of a change in mutational frequency after virus replication for several days pigs (Table 2 and Supplementary Table 2). Amino acid substitutions with a significant frequency (>2%) were found throughout all eight viral segments in clinical samples from H5N1-infected pigs; these mutations were below a frequency of 2% in the virus inoculum (Supplementary Table 2). NGS analysis revealed that oropharyngeal swab #517 on 1 DPC had the mammalian-like K526R mutation of PB2 at a frequency of approximately 3% (Table 2). Interestingly, the critical mammalian adaptation mutations, PB2-E627K and PB2-E627V, were identified in the oropharyngeal swab of pig #571 (E627K; 1 DPC), the BALF sample of pig #839 (E627K; 3 DPC), in the oropharyngeal swab of pig #94 (E627V; 1 DPC) and the BALF sample of pig #213 (E627V; 5 DPC) at a frequency of approximately 5%, 5%, 7% or 2%, respectively (see Table 2; Supplementary Table 1). Furthermore, the oropharyngeal swab of pig #94 (1 DPC) had the Q222L adaptation mutation in the HA gene at the frequency of 2.1%.

**Table 2.** Mammalian-like adaptive mutations after replication of a mink-derived clade 2.3.4.4b H5N1 in pigs.

| Viral protein | Amino acid substitution | Inoculum | Nasal swab, #517, 1 DPC | Nasal swab, #517, 3 DPC | OP swab, #94, 1 DPC | OP swab, #517, 1 DPC | OP swab, #571, 1 DPC | OP swab, #839, 1 DPC | OP swab, #634, 3 DPC | BALF, #517, 3 DPC | BALF, #575, 3 DPC | BALF, #839, 3 DPC | BALF, #94, 5 DPC | BALF, #213, 5 DPC | BALF, #714, 5 DPC |
| --- | --- | --- | --- | --- | --- | --- | --- | --- | --- | --- | --- | --- | --- | --- | --- |
| PB2 <sup>a</sup> | K526R | < 2% | < 2% | < 2% | < 2% | **3.14%** | < 2% | < 2% | < 2% | < 2% | < 2% | < 2% | < 2% | < 2% | < 2% |
|  | E627K | < 2% | < 2% | < 2% | < 2% | < 2% | **4.71%** | < 2% | < 2% | < 2% | < 2% | **5.19%** | < 2% | < 2% | < 2% |
|  | E627V | < 2% | < 2% | < 2% | **6.96%** | < 2% | < 2% | < 2% | < 2% | < 2% | < 2% | < 2% | < 2% | **2.02%** | < 2% |
| HA | Q222L | < 2% | < 2% | < 2% | **2.10%** | < 2% | < 2% | < 2% | < 2% | < 2% | < 2% | < 2% | < 2% | < 2% | < 2% |

<sup>a</sup> Amino acid substitution with a frequency >2% was not found at position of 271 of PB2 in all samples tested.

## Discussion

Rapid evolution of IAV is driven by two mechanisms: (1) the introduction of point mutations by the error-prone RNA polymerase, also called antigenic drift; and (2) by acquisition of one or more RNA segments from another influenza virus, a process called reassortment or antigenic shift. Molecular evolution allows IAVs to cross species barriers and acquire the ability to infect a wide range of hosts from avian species to swine to humans. This broad host tropism provides opportunities for IAVs to gain distinct geno- and phenotypes, to attain pandemic potential, and to evade pre-existing immunity in humans ^24^. Specific host-virus-environment characteristics accelerate the emergence of novel IAVs with pandemic potential. For pandemic preparedness, it is critical to assess the potential pandemic risk of newly emerged IAVs; the evaluation of certain virus characteristics is mainly done by *in vitro* and sometimes also *in vivo* experiments ^25^. However, it is essential to assess the pathogenesis and transmissibility of IAVs in target animal models. Pigs are natural hosts for IAVs, and it is known that certain IAVs can be efficiently transmitted between pigs. Importantly, interspecies transmissions between human and pigs and *vice versa* are not uncommon and are facilitated by an extensive human-pig interface in various agricultural settings^26^. Furthermore, pigs express receptors specific for both, mammalian and avian IAVs, providing opportunities for co-infections with these viruses and the subsequent generation of novel reassortant influenza viruses ^27^. Recently, interspecies transmission of clade 2.3.4.4b H5N1 virus to mink and sustained mink-to-mink transmission was reported in Spain, posing a significant concern to public health^13^. Therefore, it was crucial to evaluate the pathogenicity and transmissibility of the recently emerged clade 2.3.4.4b H5N1 virus isolated from diseased mink in domestic pigs.

One of key findings in this study was that experimental infection of pigs with the mink-derived clade 2.3.4.4b H5N1 virus resulted in productive virus replication and seroconversion in 100% of the principal-infected pigs. Virus titers in BALF samples of principal-infected animals reached 10^3.3^ to 10^4.7^ TCID_50_/mL at 3 DPC and declined to 10^2.7^ to 10^3.5^ TCID_50_/mL at 5 DPC. In addition, the virus was found in other respiratory tissues, such as trachea, bronchi, as well as lymphoid tissues, such as tracheobronchial lymph nodes and tonsils. Moreover, gross and histological evaluation of various tissues demonstrated that the infection caused acute multifocal to coalescing necrotizing broncho-interstitial pneumonia at 3 and 5 DPC, with residual mild interstitial pneumonia still present at 21 DPC. Based on the microscopic lesion distribution, virus titers and viral antigen/RNA tissue expression, it is clear that the lesions and distribution of this mink-derived clade 2.3.4.4b H5N1 virus mostly involved the lower respiratory tract (i.e., pulmonary parenchyma) with only mild alterations on the upper respiratory tract. Previous studies demonstrated virus replication of avian-derived H5Nx clade 2.3.4.4 HPAI viruses from North America in the lower respiratory tract of pigs upon experimental infection ^28^. In the latter study, infection of pigs with the avian-origin H5Nx clade 2.3.4.4 viruses, isolated in 2014 and 2015 in the U.S, led to viral RNA detection in 80-100% of the animals and successful virus isolation in 60-100% of the BALF samples at 3 and 5 DPC ^28^. Consistent with virus detection in the majority of animals, 60-100% of infected animals seroconverted by 21 DPC (28). In addition, in a very recent study, low susceptibility of pigs against experimental infection with an avian-derived H5N1 clade 2.3.4.4b virus, isolated from chickens in Germany in 2022, was reported ^29^. This chicken H5N1 clade 2.3.4.4b isolate lacked any mammalian-adaptive mutations. Nasal and alimentary exposure of pigs to this avian-derived H5N21 clade 2.3.4.4b virus only resulted in marginal virus replication, without inducing any clinical signs or pathological changes. Only 1 out of 8 pigs seroconverted, indicating a rather high resistance of pigs to infection with the avian-derived derived H5N21 clade 2.3.4.4b virus without mammalian-like adaptations ^(29)^.

Experimental infection of pigs with LPAI viruses without mammalian-like adaptations were also previously performed, but none of the experimental infections with LPAI H5 viruses resulted in a productive infection ^30^. We conclude from our present study with the mink-derived clade 2.3.4.4b H5N1 HPAI virus, that this virus is not only able to efficiently infect mink, but also pigs. The mink-derived H5N1 clade 2.3.4.4b virus efficiently replicated in the lower respiratory tract of pigs inducing sizeable lung lesions; in addition, nasal and oral shedding was observed on early days after infection although less efficient than reported for swine-derived influenza viruses ^31,32^. However, it is important to note that the mink-derived H5N1 clade 2.3.4.4b virus was unable to spread from principal-infected to co-mingled in-contact sentinel animals; this was most likely due to the limited amount of virus shedding by the nasal and oral cavities in principal-infected pigs. Interestingly, neither infectious virus, viral RNA nor histopathological changes were present in tissues from the central nervous system of principal-infected pigs in our study. In recent spillover events, wild mammals infected with the clade 2.3.4.4b H5N1 virus primarily exhibited neurological symptoms with histopathological changes in brains as well as viral RNA detection in CNS tissues ^33-35^. Similarly, the mink infected with the H5N1 clade 2.3.4.4b virus used in this study also showed neurological manifestations, such as ataxia and tremors ^13^. We could not see any neurological clinical signs in principal-infected pigs, indicating that the pathogenicity of the H5N1 clade 2.3.4.4b viruses could vary in different mammalian host species.

Earlier studies showed that experimental infection of pigs with avian-origin LPAI H5 subtype viruses isolated between 1980 and 2005 led to nasal shedding of infectious virus of up to 10^5.5^ egg infectious doses (EID)_50_/mL and 10^5.3^ EID_50_/100 mg ^36,37^. In addition, it was shown that experimental infection of pigs with human and avian-origin HPAI H5N1 viruses isolated in early (1997) and later (2004) in the emergence of the HPAIV H5N1 resulted in the isolation of infectious virus in nasal swabs/nasal tracts over the course of infection, but no pig-to-pig transmission ^38,39^. When pigs were infected with the index human H5N1 HPAIV strain (HK156-97) and an initial chicken H5N1 HPAIV isolate (CHK258-97) by the oral and nasal routes, both the human and chicken H5N1 viruses replicated well in the nasal tract of pigs, with the chicken isolate to higher titers than the human virus. Importantly, neither the human nor the chicken H5N1 virus transmitted to contact pigs in the same pen. Thus, pigs support the replication of the index human and an early chicken H5N1 virus to modest titers, and there was no detectable transmission to contact animals ^38^. Similarly, experimental studies on the replication and transmissibility of 2004 avian H5N1 HPAI viruses associated with human infections in Vietnam to pigs revealed that H5N1 all viruses tested replicated in the swine respiratory tract but none were transmitted to contact pigs (39).

Epidemiological studies showed limited serologic evidence (8/3175 pigs positive; 0.25%) of exposure to the HPAI H5N1 viruses in Vietnamese pigs ^39^. These findings indicate that pigs can be infected with the 2004 Vietnamese HPAI H5N1 viruses but that these viruses are not readily transmitted between pigs under experimental conditions ^39^. In contrast to the above described work, no viral RNA was detected in nasal swabs of pigs infected with avian-derived clade 2.3.4.4 H5Nx viruses in 2014 ^28^, and only limited detection of viral RNA but failure to isolate infectious virus was reported when pigs were infected with an avian-derived clade 2.3.4.4b H5N1 virus without mammalian-like adaptations ^29^. Collectively, these results suggest that some isolates of HPAI H5 subtype viruses can replicate efficiently in the upper respiratory tract of pigs, with limited replication in the lower respiratory tract. There is no transmission of these viruses to co-mingled contact sentinels.

Through the continuous evolution of the HPAI H5Nx viruses over two decades, the A/Goose/Guangdong/1/1996 H5N1 lineage (GsGD) may have acquired increased fitness in avian species resulting in efficient spread across the globe. However, the continued adaptation of the H5N1 clade 2.3.4.4b virus to avian species may, in turn, contribute to a loss of fitness for mammalian species, which resulted in its reduced replication capacity in the upper respiratory tract of pigs. Despite the shedding of infectious virus in nasal and oropharyngeal secretions from principal-infected pigs, there was no evidence of transmission of the mink-derived HPAI H5N1 clade 2.3.4.4b virus to sentinel pigs under our experimental conditions. This result was consistent with previous findings indicating a lack of pig-to-pig transmission for H5- or H7-subtype HPAIVs, regardless of the presence of infectious virus in nasal and/or oral excretions ^28,29,38,39^. The lack of transmissibility of the mink-derived H5N1 clade 2.3.4.4b isolate in pigs is supported by the absence of seroconversion in the co-mingled sentinel animal; also, this virus did not transmitted to the 11 farm workers that had been in close contact with the H5N1-infected minks at the site of the outbreak ^13^. Furthermore, a recent study showed that two other mink-derived H5N1 viruses which differ by several amino acids in the PB2, PB1, PA, NA, NS1 and NS2 proteins, did not support transmission to contact animals (neighboring cages) through respiratory droplets in the ferret model ^40^. Importantly, it has been previously documented that H5 subtype viruses can acquire the ability to transmit efficiently among pigs through reassortment. Lee *et al.* ^41^ isolated two H5N2 viruses from pigs showing typical clinical symptoms of influenza-like illness; one was an avian-origin virus, the other one was an avian-swine reassortant virus with PB2, PA, NP and M genes derived from swine influenza viruses. Upon experimental infection of pigs, both viruses could be isolated from nasal swabs, but pig-to-pig transmission occurred only in the group infected with the reassortant H5N2 virus ^41^.

One of the important findings is that the H5N1 viruses inoculated into pigs acquired mammalian-like mutations/adaptations in the course of several days of pig infection, which potentially results in increased fitness of the H5N1 virus in mammalian hosts. There is a total of four mink-derived H5N1 isolates from the infected Spanish farm which were sequenced; all of them possessed the PB2 T271A mutation. The PB2 T271A mutation is known to lead to enhanced polymerase activity in mammalian cells and might have contributed to cross-species transmission from avian species to mammals including mink ^13,42^. Our results show that the T271A mutation was stable with no evidence of a change in the frequency of the mutation after replication of the virus for several days in pigs. However, minor variants possessing the mammalian-like E627K mutation in PB2 emerged in oropharyngeal swab (pig #714) and BALF (pig #839) samples obtained from principal-infected pigs, even though they were found at low frequencies (4.7% and 5.2%, respectively). The E627K mutation is a key determinant for mammalian adaptation of avian influenza viruses; its introduction into the genome of an avian influenza virus enables efficient replication of the avian-origin polymerase complex in mammalian cells ^43^. It is well known that H5N1 viruses harboring the PB2 E627K mutation have increased virulence in mice; it also contributes to air-borne virus transmission in ferrets and contact transmission in guinea pigs ^44-48^. Similarly, an amino acid substitution from glutamic acid (E) to valine (V) at position 627 of the PB2 gene (E627V) was shown to increase viral replication in mammalian cells and virulence in mice models ^49^; an E627V mutation was detected as a minor virus population in the oropharyngeal swab of pig #94 and BALF sample of pig #213. Furthermore, the oropharyngeal swab of gig #517 possessed a PB2 K526R mutation at the frequency of 3.1%; the K526R mutation was shown to enhance the effect of the PB2 E627K on avian influenza virus replication in mammalian cells and also their virulence in mice ^50^. Another key mutation is the Q222L (according to the H5 numbering system) in the HA gene which is equivalent to a Q226L mutation using the H3 numbering system. This mutation was found in oropharyngeal swab of pig #94 at 1 DPC. Amino acid (AA) position 222 in the HA is involved in the binding of HA to its sialic acid receptor and plays a critical role in determining the host range of IAVs. The HA of mammalian IAVs binds to α2,6-linked sialic acid, whereas the HA of avian IAVs binds to α2,3-linked sialic acid. Several amino acids in the HA receptor binding site are responsible for switching sialic acid specificity; however, the Q222L and G224S AA positions in the H5 numbering system, which are equivalent to AA positions Q226L and G228S in the H3 numbering system, are crucial for increased α2,6-sialic acid receptor preference in different subtypes of avian IAVs; they are located in the receptor binding domain and interact directly with the host’s sialic acid receptor ^51^. In addition, the Q222L mutation was previously described to contribute to the emergence of transmissible H5N1 viruses via aerosols after serial passage of H5N1 in ferrets ^45,46^. Despite the critical role of these mutations in pathogenicity and transmissibility of IAVs in mammalian hosts, it still remains unclear how the H5N1 clade 2.3.4.4b virus underwent its molecular evolution, and eventually acquired critical mammalian-like mutations. In addition to the role of pigs in the ecology of IAVs as the “mixing vessel”, the emergence of mammalian-adapted H5N1 viruses in pigs is close to reality; therefore, pigs may serve as the “hotspot” to introduce mammalian-like mutations into avian influenza viruses although the frequencies of the mammalian-like mutations described here are rather low.

Spillover of H5N1 HPAI clade 2.3.4.4b viruses to mammals and their sustained transmission between mammals has been reported ^13,52^. The majority of mammals infected by the HPAI H5N1 clade 2.3.4.4b virus are carnivores, and the consumption of infected dead wild birds is presumably the basis for these spillover events ^53^. In contrast to the HPAI H5N1 clade 2.3.4.4b virus infections in wild avians and wild mammals, outbreaks in farmed avian and mammal species pose a much greater risk to humans, due to its close proximity to occupational workers and -in the case of mammalian species - the favorable environment for the acquisition of mammalian-like adaptations. Therefore, evaluating the infectivity/susceptibility and transmissibility of the emerging HPAI viruses in well-established mammalian animal models such as swine and ferrets is crucial to determine their pandemic potential. The present study demonstrated that infection of pigs with the mink-derived clade 2.3.4.4b H5N1 virus led to a productive viral replication in the respiratory tracts of all principal-infected pigs including seroconversion; however, virus shedding from principal-infected pigs was not high and/or frequent enough for an efficient transmission to co-mingled sentinel pigs. Notably, key mammalian-like mutations were found in clinical and/or tissue samples derived from principal-infected pigs. In conclusion, the results obtained in the present study suggest that the mink-derived H5N1 clade 2.3.4.4b virus exhibited increased infectivity in pigs when compared to avian-origin H5N1 clade 2.3.4.4b viruses, however, did not transmit to co-mingled sentinel animals; therefore, this virus would still be placed in a moderate risk group in terms of transmission ability to humans.

## Acknowledgments

We gratefully thank Yonghai Li, Patricia Assato, Michelle Zajac, Isaac Fitz for technical assistance, and Innovative Diagnostics for providing ELISA kits. Funding for this study was provided through grants from the NIAID supported Center of Excellence for Influenza Research and Response (CEIRR) under contract number 75N93021C00016, the National Bio and Agro-Defense Facility (NBAF) Transition Fund from the State of Kansas, the MCB Core of the Center on Emerging and Zoonotic Infectious Diseases (CEZID) of the National Institutes of General Medical Sciences under award number P20GM130448, the NIAID Centers of Excellence for Influenza Research and Surveillance (CEIRS) under contract number HHSN 272201400006C.

## Author contributions

J.A.R and N.O. conceptualized the research project. T.K., J.D.T., M.C., E.L.L., C.D.M., K.C., F.S.M., T.J., I.M., N.N.G., U.B.R.B., N.O., R.J.W. and J.A.R acquired the data.

---

**Supplementary Figure 1.**
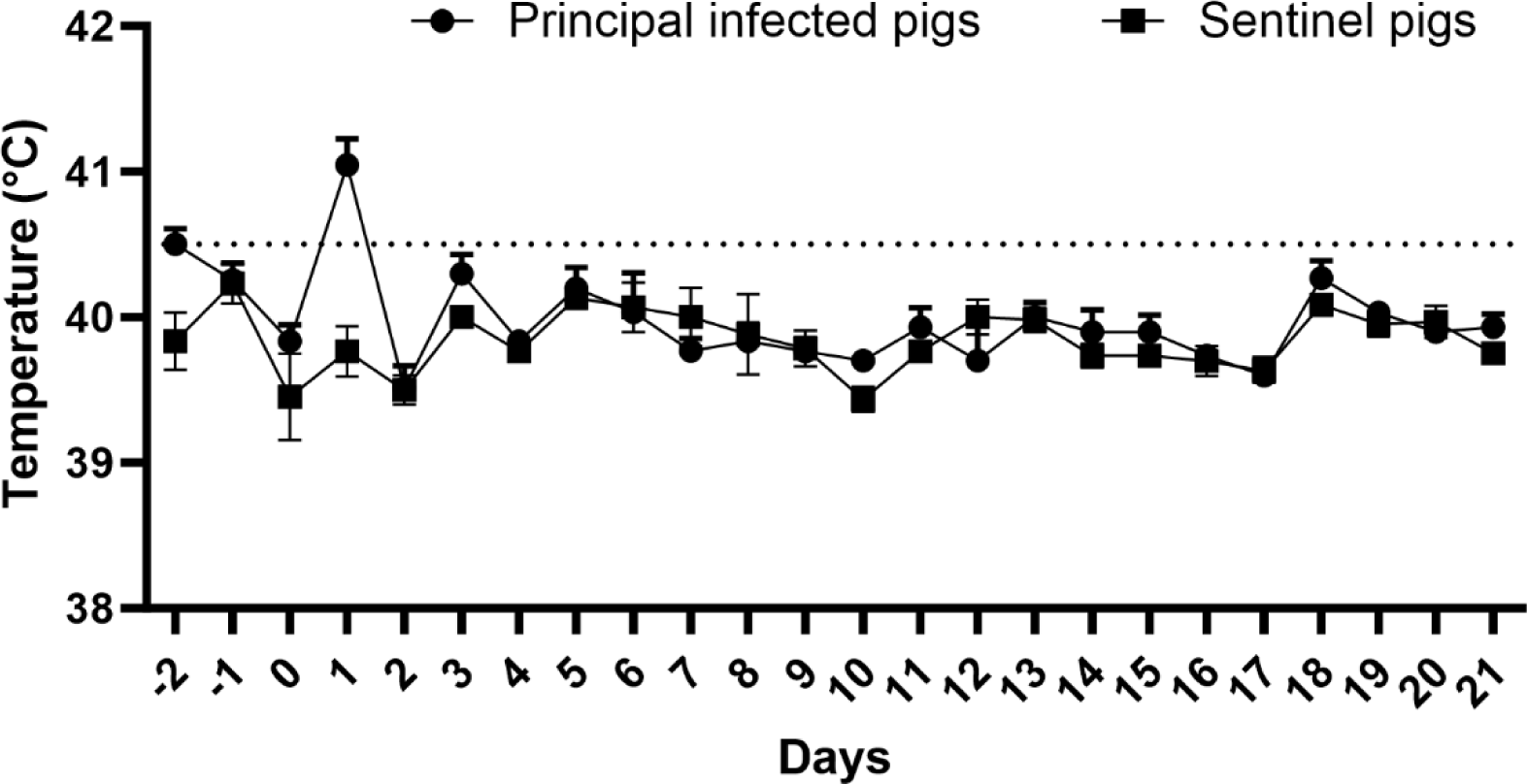
Daily body temperature of principal infected and sentinels in this study. The value is expressed as the mean ± SEM. The dash line indicates 40.5 °C, which is considered fever in this study.

**Supplementary Figure 2.**
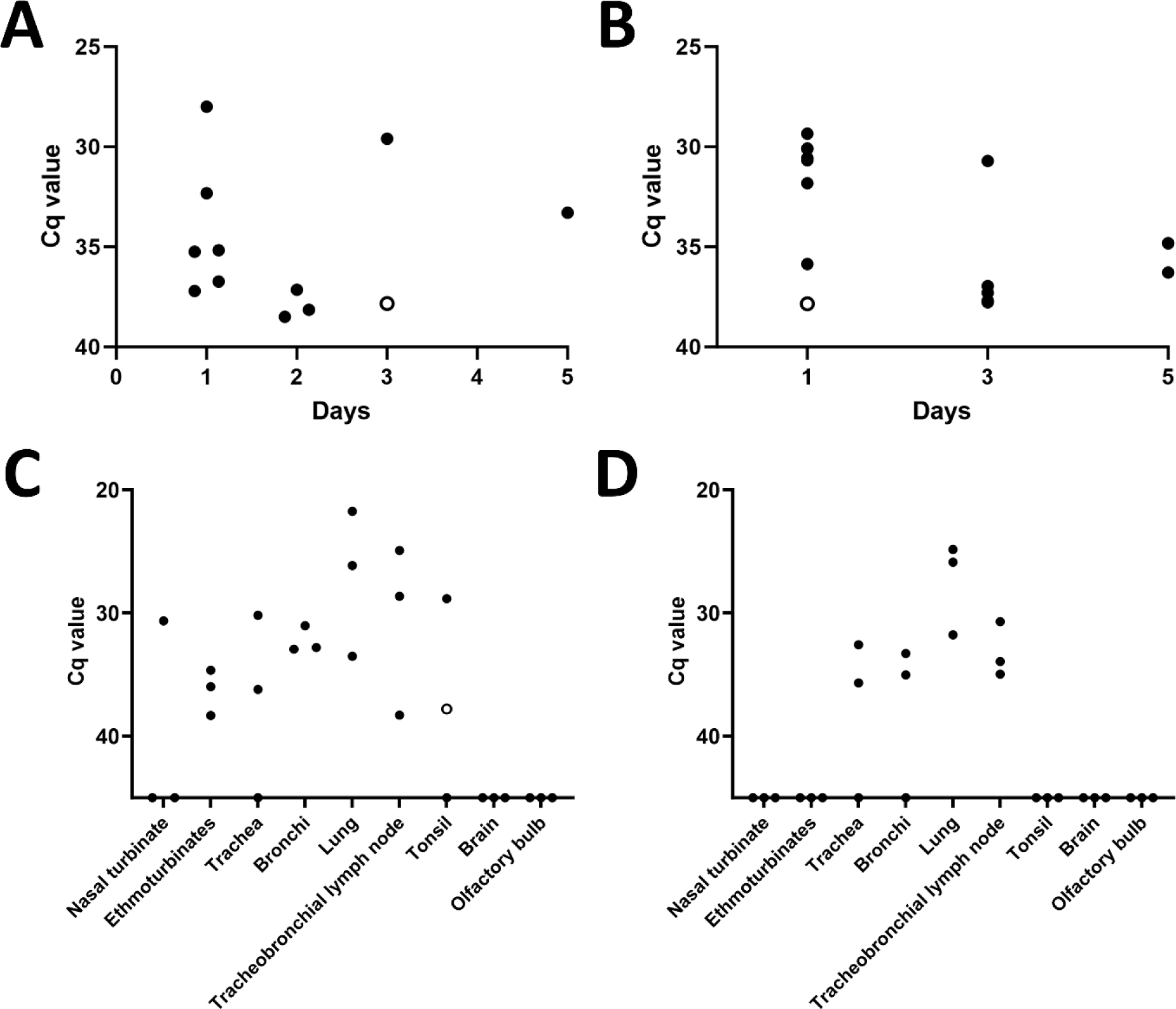
Viral RNA detection in (A) nasal swab and (B) oropharyngeal swab of H5N1-infected pigs (n=9, at 1, 2, and 3 days post-challenge (DPC); and n=6, at 4 and 5 DPC). The Cq values of positive swabs were shown in the graphs. The distribution of viral RNA in tissues of H5N1-infected pigs at (C) 3 DPC and (D) 5 DPC. Empty circles represent the single well positive in duplicates.

**Supplementary Figure 3.**
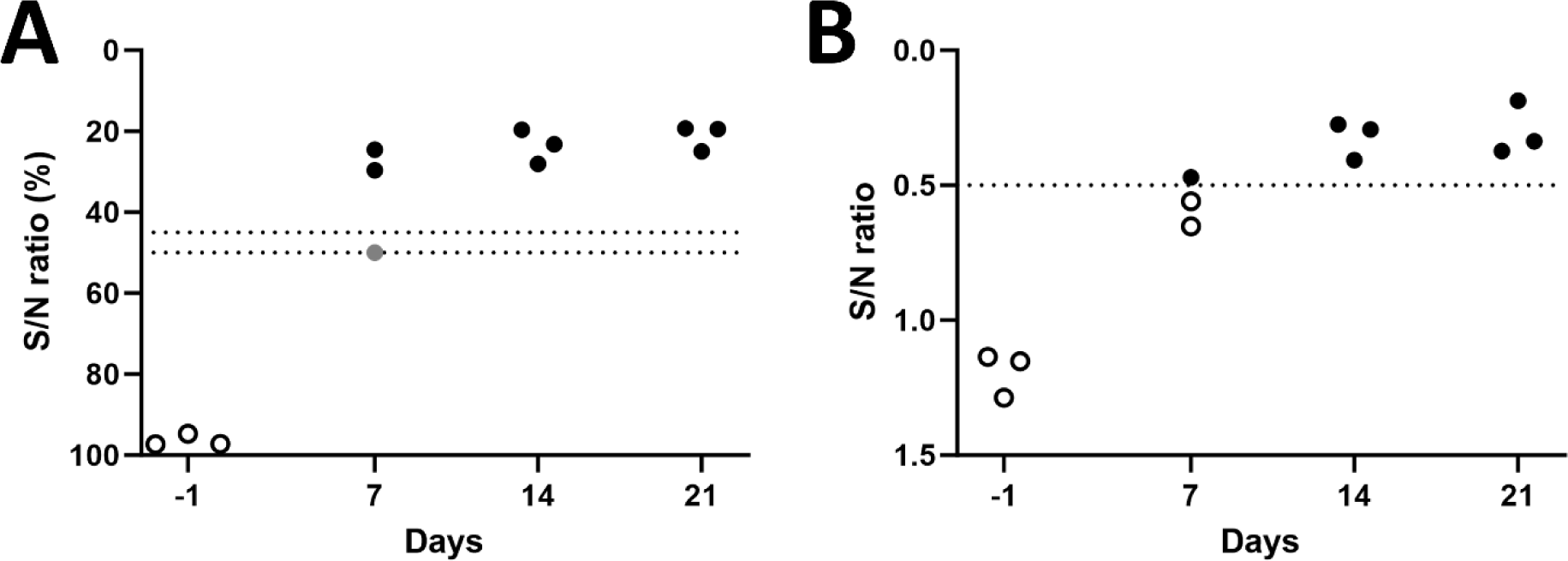
Antibody responses in H5N1-infected pigs employing two competitive ELISAs: (A) ID Screen® Influenza A Antibody Competition Multi-Species and (B) IDEXX AI MultiS-Screen Ab Test. Empty and grey circles represent negative and suspicious.

**Supplementary Figure 4.**
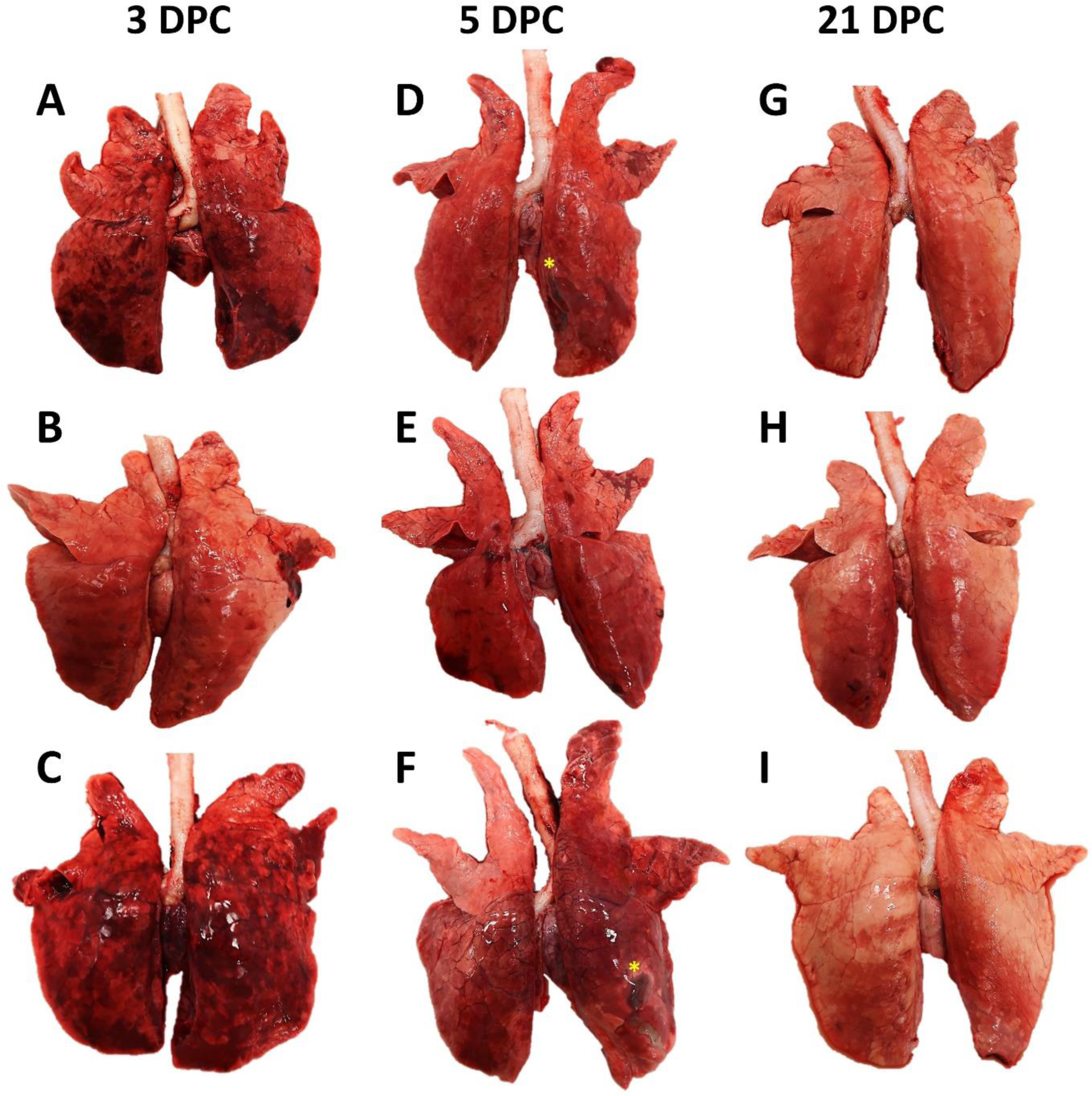
Gross changes in the lungs of a mink-derived clade 2.3.4.4b H5N1 virus-infected pigs at 3, 5 and 21 DPC. At 3 DPC, two pigs (A and B) had moderate to severe, multifocal to coalescing interstitial pneumonia with congestion and edema affecting all lobes (A and B), while the third animal (C) had mild, multifocal pneumonia with congestion affecting predominately the right lung and left caudal lung lobes (C). At 5 DPC (D–F), coalescing areas of pulmonary consolidation with congestion and edema were noted in all pigs. The right middle and caudal lobes were most commonly affected (D–F) with occasional bullae (asterisk). At 21 DPC, only a few areas of pulmonary consolidation were noted (G–I).

**Supplementary Figure 5.**
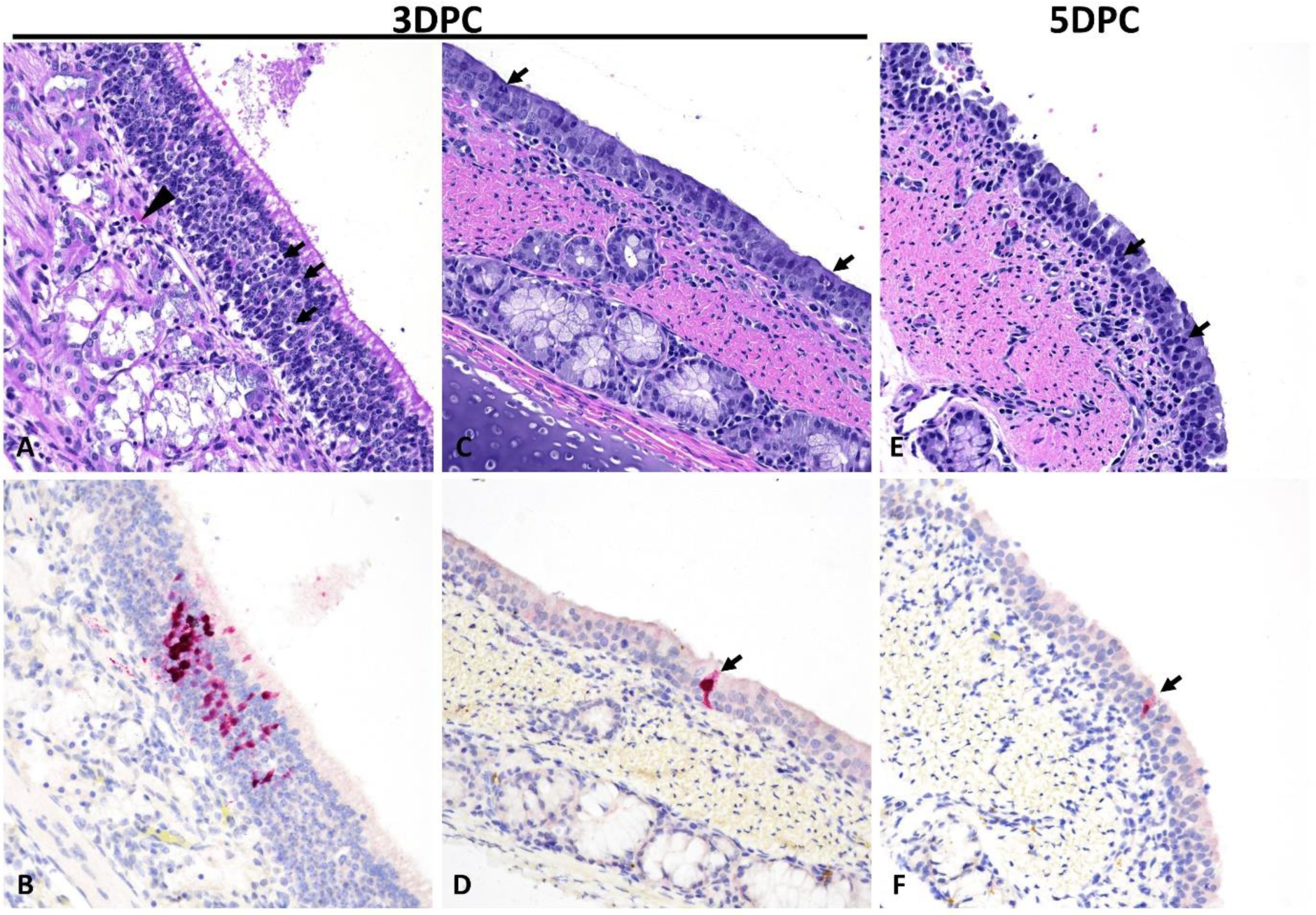
Histological and immunohistochemical findings in the upper respiratory tract of pigs infected with a mink-derived clade 2.3.4.4b H5N1 virus. In a single animal, a focal area of the olfactory epithelium is mildly infiltrated by few lymphocytes (A, arrows) that also mildly infiltrate the subjacent lamina propria (A, arrowhead) at 3 DPC. Within this area, olfactory epithelia contain abundant influenza NP antigen (B, intracytoplasmic and intranuclear). At 3 and 5 DPC, there is mild lymphohistiocytic and neutrophilic tracheitis (C and E) with lymphocytes and histiocytes within the lamina propria and sporadic neutrophils and lymphocytes transmigrating through the lining epithelium (arrows). Sporadic epithelial cells contain intracytoplasmic and intranuclear influenza NP antigen (D and F, arrows). H&E (A, C, E) and IHC (Fast Red, B, D and F), total magnification 200X.

**Supplementary Figure 6.**
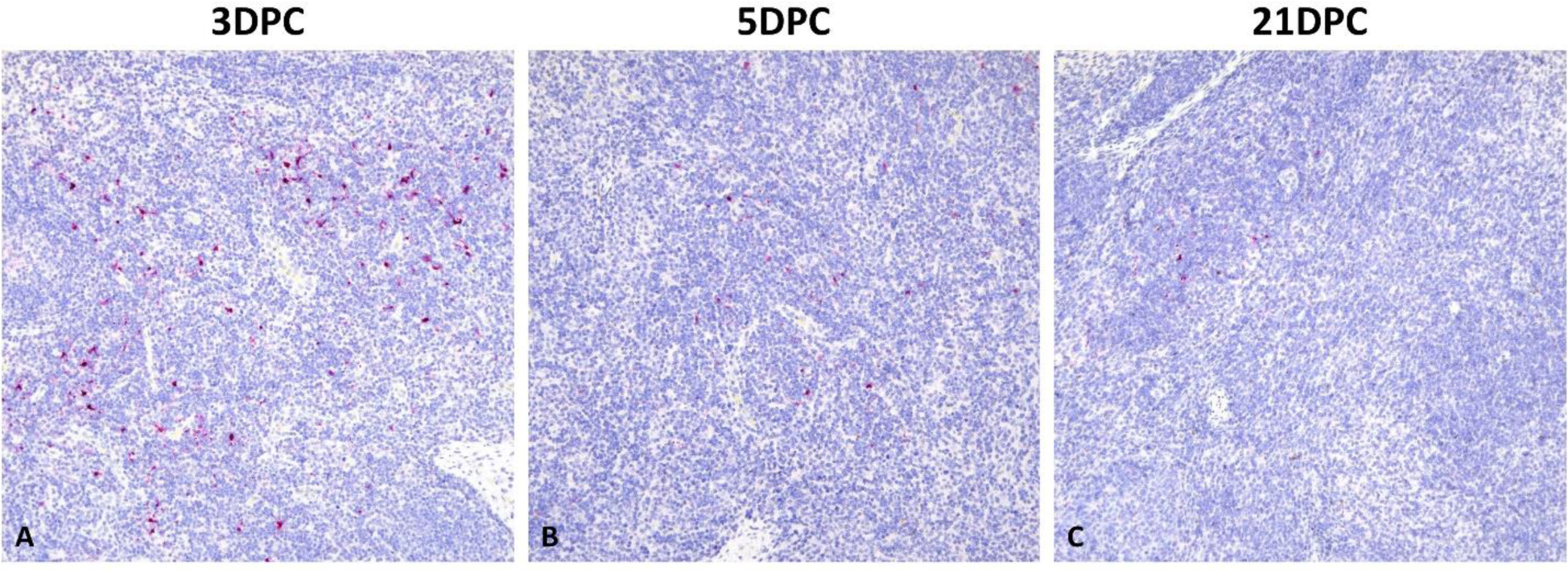
Immunohistochemical findings in the tracheobronchial lymph nodes of pigs infected with a mink-derived clade 2.3.4.4b H5N1 virus. At 3 DPC (A), there are numerous influenza NP antigen-positive cells with dendritic projections of their cytoplasm (consistent with antigen presenting cells). The number of immune-positive cells gradually declines at 5 and 21 DPC (B and C, respectively). IHC, Fast Red, total magnification 100X.

**Supplementary Table 2.**
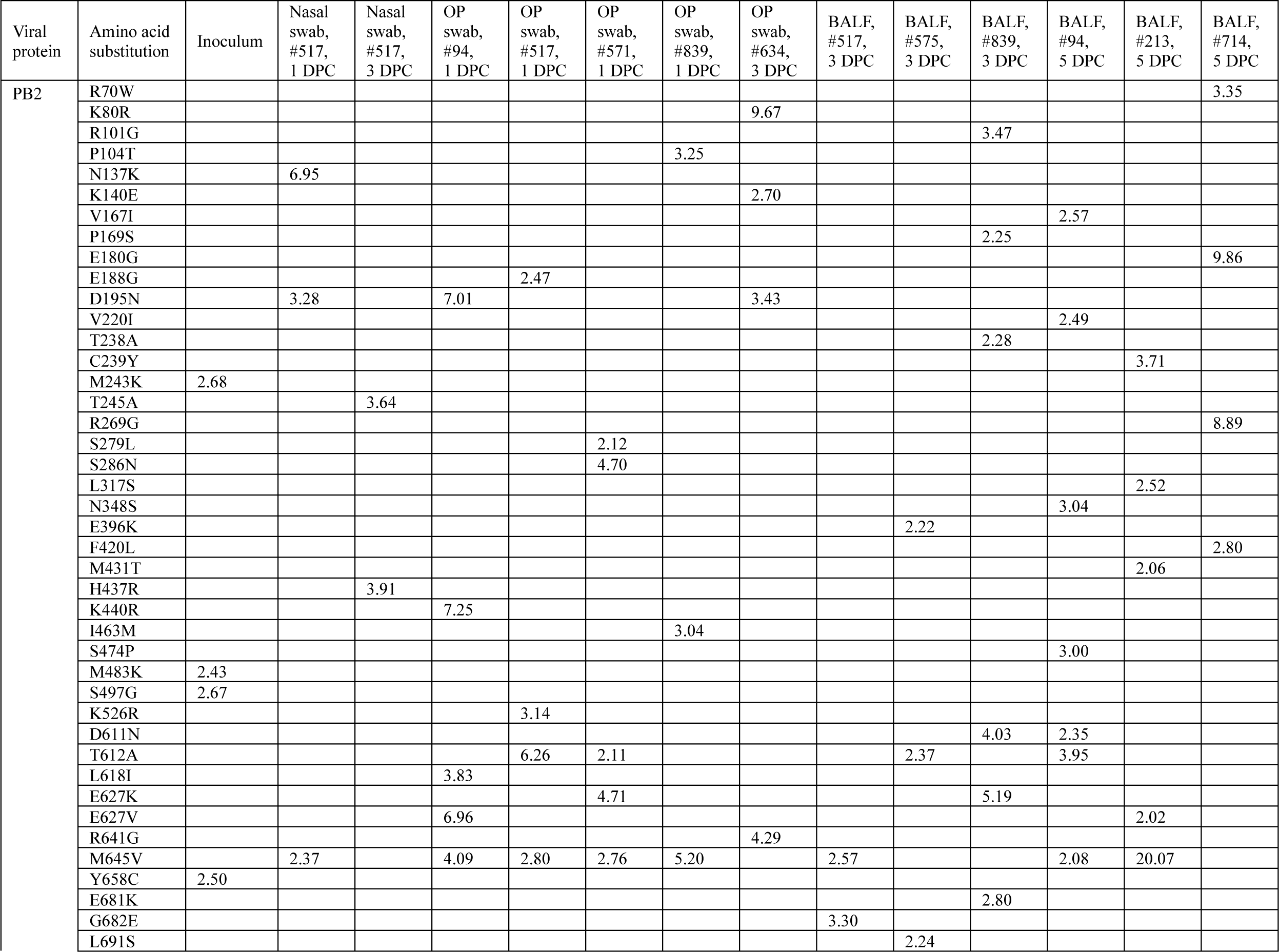
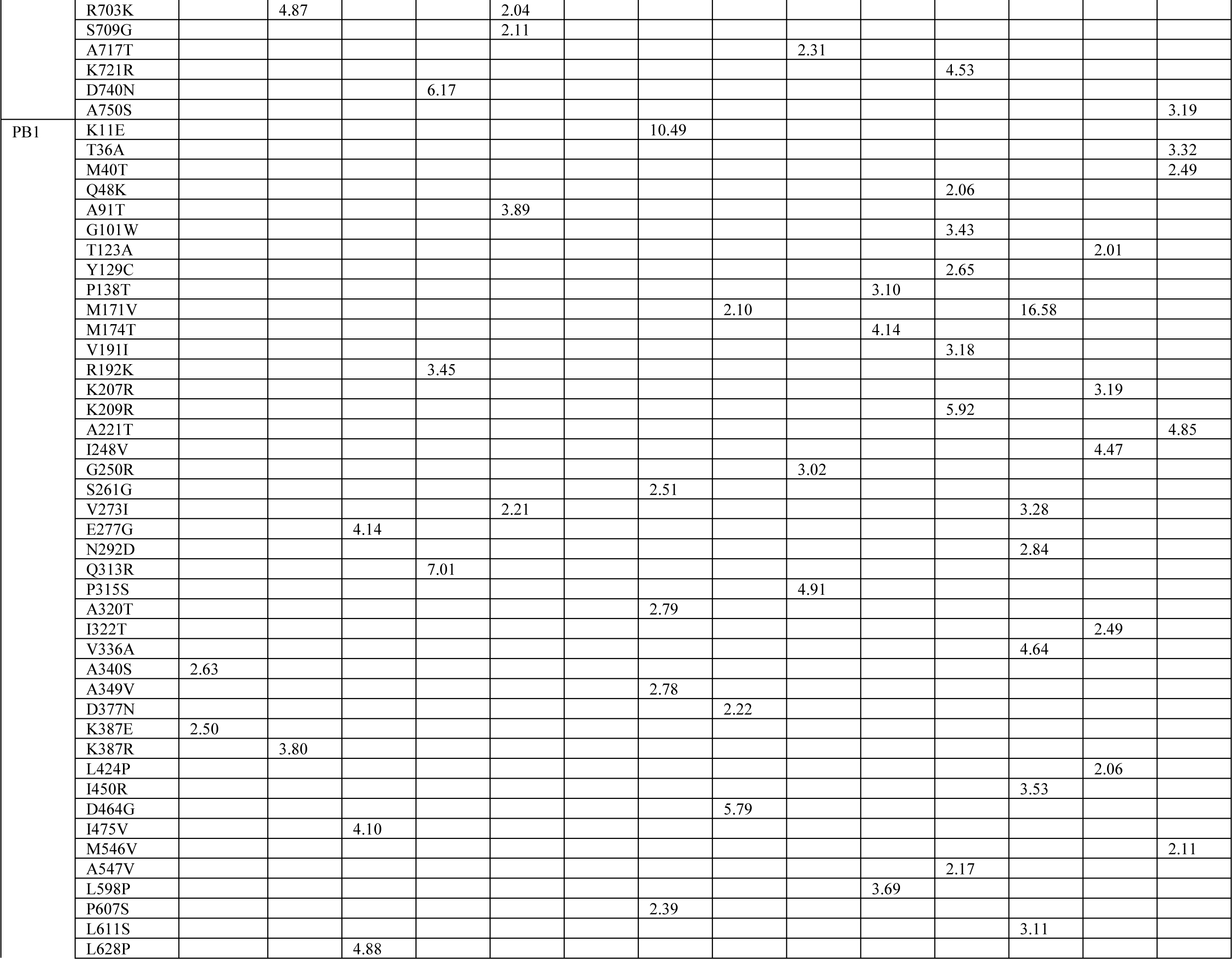
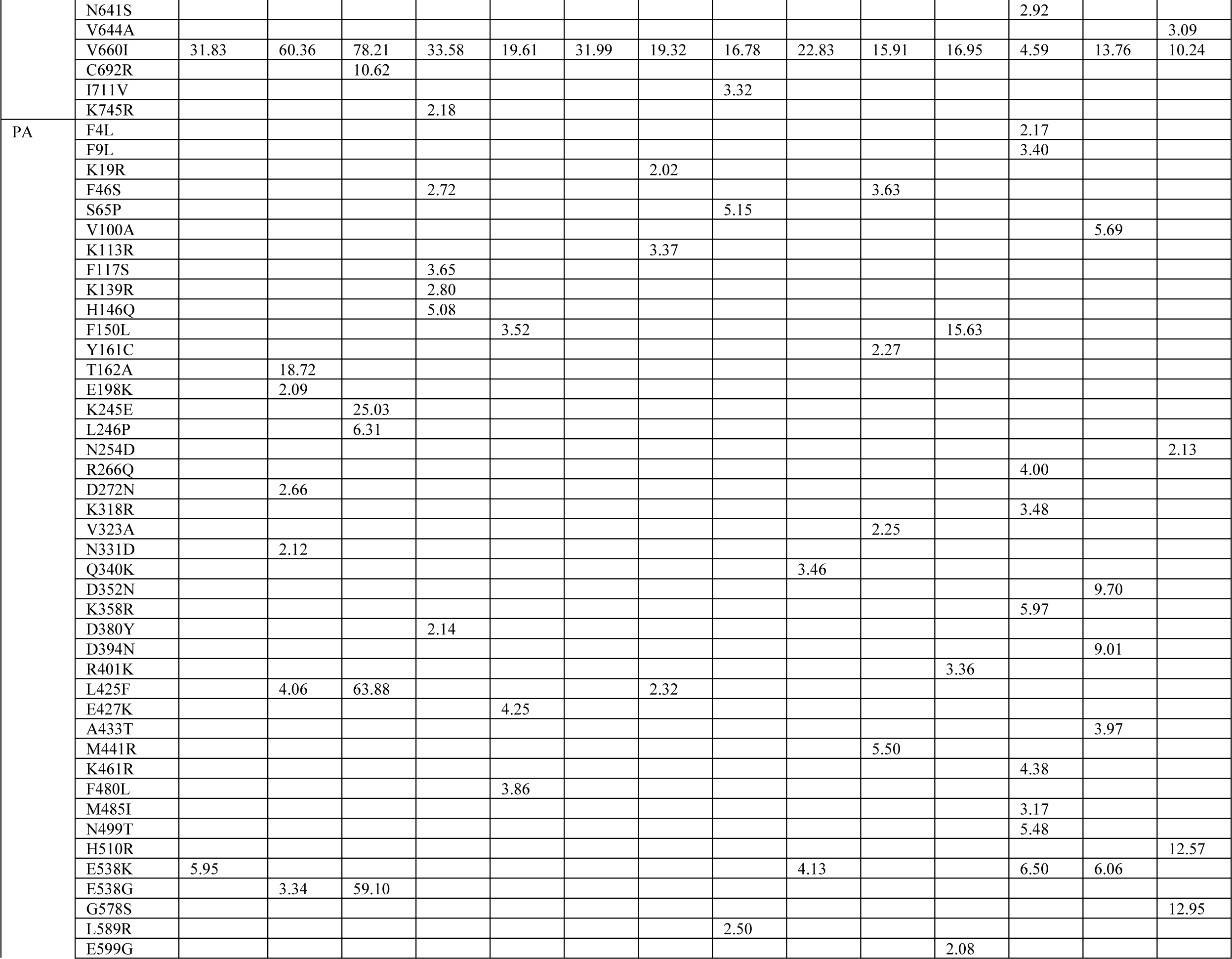
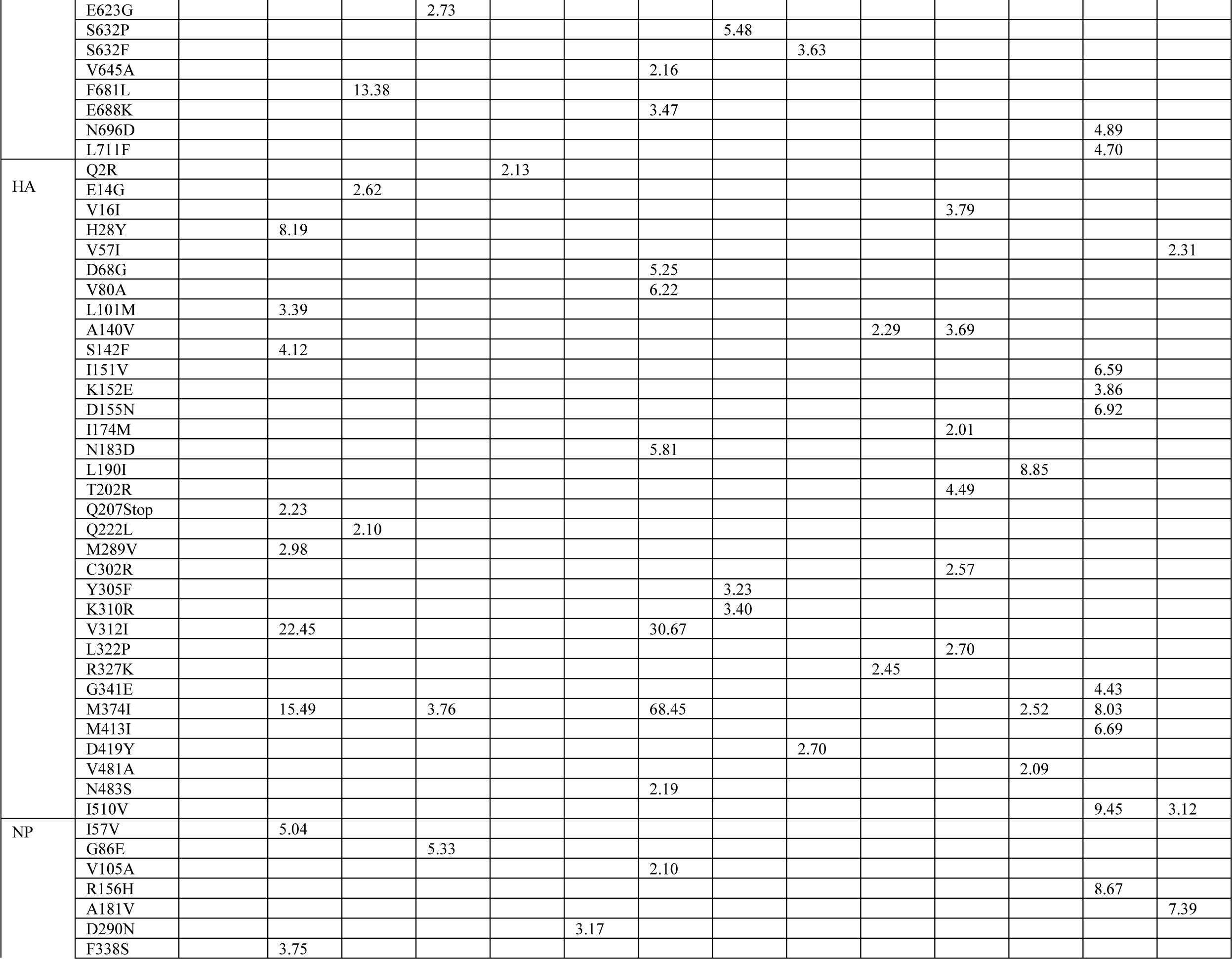
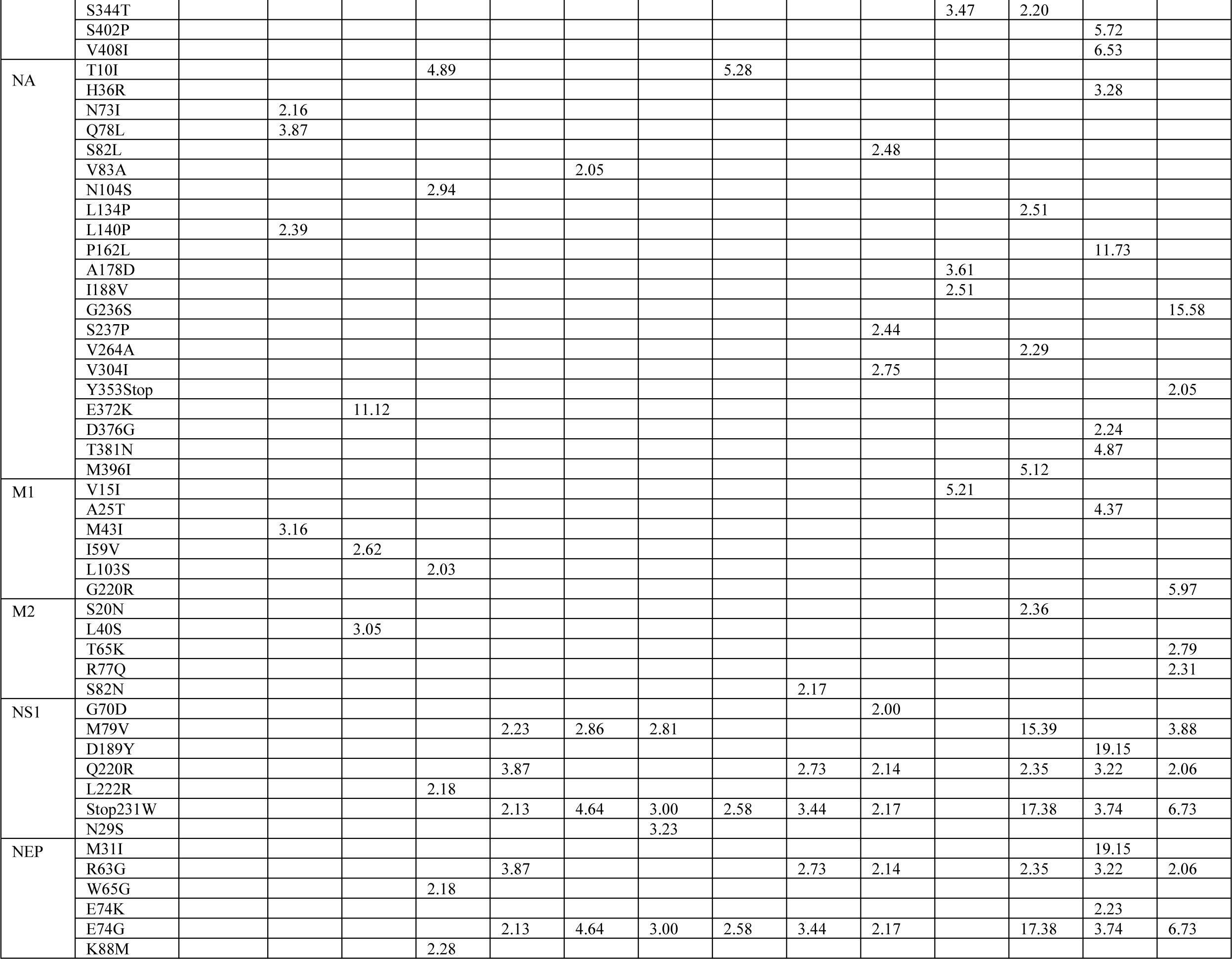
Frequencies of mutations of a mink-derived clade 2.3.4.4b H5N1 in pigs. Blank represents less than 2%.

